# Generating Synthetic MR Perfusion Maps from DWI and FLAIR in Acute Ischemic Stroke: Development and External Validation of a Deep Learning Model

**DOI:** 10.1101/2025.10.23.684079

**Authors:** Anna Matsulevits, Alexander Koch, Clara Mahé-Verdure, Martin Bendszus, Adam Hilbert, Mary Boullet, Gaultier Marnat, Matthias A. Mutke, Orhun Utku Aydin, Stéphane Olindo, Igor Sibon, Michel Thiebaut de Schotten, Dietmar Frey, Thomas Tourdias

## Abstract

**Background:** Magnetic resonance imaging (MRI) is critical for acute stroke triage, but time-consuming, and often requires contrast injection for perfusion imaging. This study aimed to synthesize T-map perfusion maps from routinely available, non-contrast DWI and FLAIR using deep generative models. We hypothesized that relevant perfusion information could be inferred from these modalities to streamline imaging and reduce reliance on dynamic susceptibility contrast perfusion.

**Methods:** Acute MRI data from 355 patients with anterior circulation stroke, including dynamic susceptibility contrast perfusion, were retrospectively collected from two European centers (Heidelberg: 2010–2018; Bordeaux: 2021–2022). Six versions of a denoising diffusion probabilistic model (DDPM) and a GAN architecture were trained to generate synthetic T-max perfusion maps from DWI, FLAIR, and infarct core mask as inputs. Performance was assessed by comparing synthetic and ground truth T-max maps using image similarity metrics. Regions with T-max >6s were compared using Dice coefficients, and mismatch volume distributions were analyzed. An ablation study quantified the contribution of each input.

**Results:** The best performance was achieved by a DDPM with a 2.5D architecture using DWI, FLAIR, infarct core mask, and a perfusion-weighted loss function. It produced synthetic perfusion T-max maps with high similarity to ground truth under 110 seconds. The model showed strong spatial overlap for T-max >6s regions in internal validation (average Dice = 0.82, SD = 0.08), and external validation average (Dice 0.59, SD = 0.13), respectively. Synthetic maps closely matched ground-truth mismatch distributions, capturing key perfusion patterns. The infarct core mask played a critical role in model performance, alongside DWI and FLAIR inputs.

**Conclusions:** We propose a non-invasive, scalable framework to generate synthetic T-max perfusion maps from non-contrast MRI. This approach could expand access to perfusion data in acute stroke, shorten imaging protocols, and accelerate treatment decisions by eliminating the need for contrast-enhanced acquisition.

**Graphical abstract:** 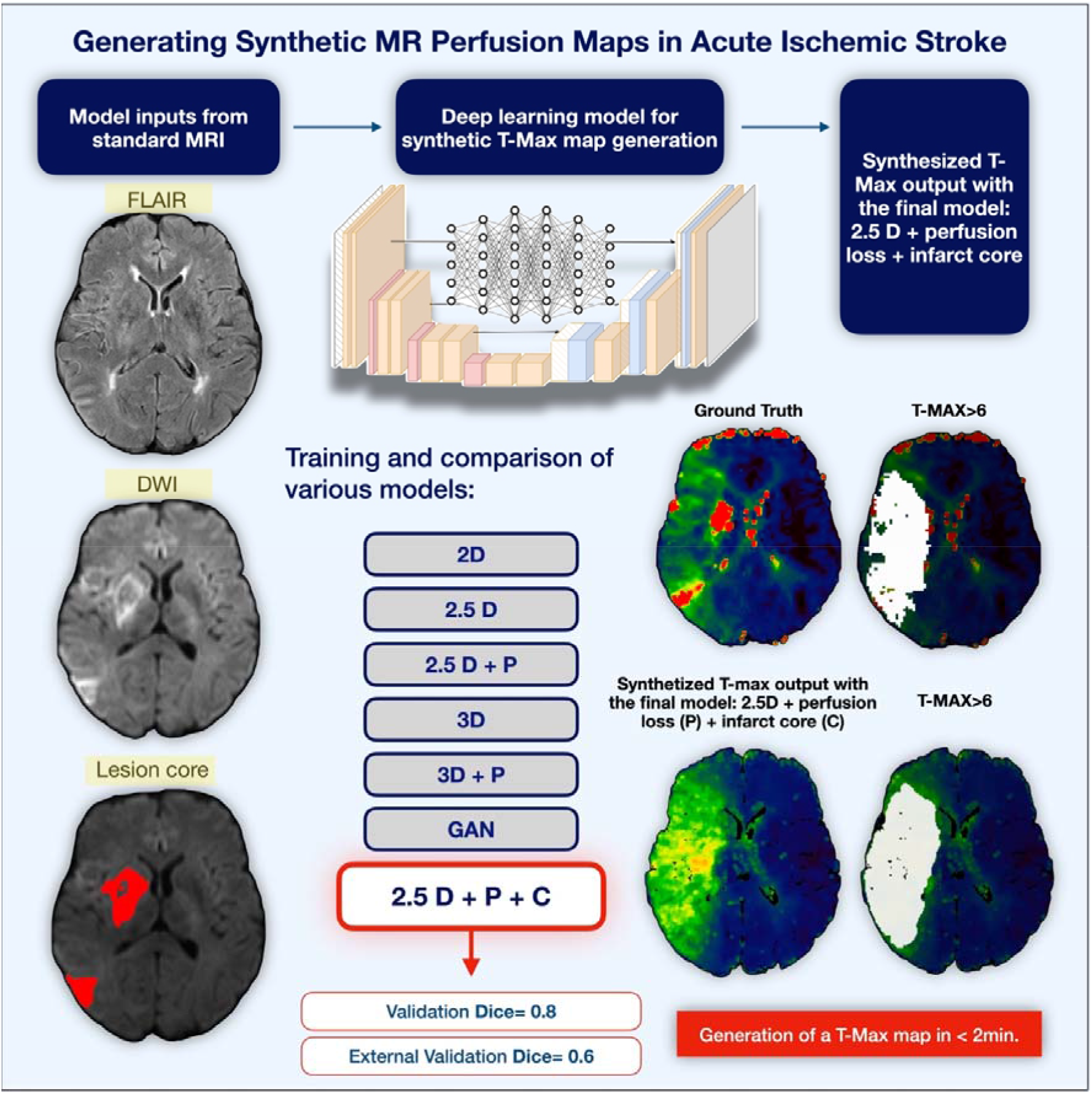

## Introduction

The effective management of ischemic stroke hinges on revascularization strategies that must be implemented as quickly as possible to salvage viable brain tissue. Consequently, the triage of acute ischemic stroke patients relies heavily on the rapid acquisition and interpretation of neuroimaging.

While computed tomography (CT) is the most widely used imaging modality worldwide, acute magnetic resonance imaging (MRI) offers specific advantages. These include (i) a substantial reduction in intravenous thrombolysis for stroke mimics ^1^, *(ii)* the ability to assess eligibility for thrombolysis in cases of unknown symptom onset^2^, and *(iii)* and a significant decrease in the need for repeated imaging during the course of hospitalization^1^, and lastly a radiation-free assessment. With MRI scanners now more commonly located within emergency departments and most metal implants proven to be MRI-safe^3^, a growing number of comprehensive stroke centers may consider MRI as the first-line imaging modality for acute stroke evaluation^1^. However, MRI has a longer acquisition time (approximately 13 minutes compared to 9 minutes for CT^4^), which can be associated with a pejorative workflow delay, even though this difference does not seem to adversely affect functional outcomes^4^.

In this evolving context, efforts to streamline MRI protocols and reduce scan time without compromising diagnostic value are of increasing relevance. A standard acute stroke MRI typically includes diffusion-weighted imaging (DWI), fluid-attenuated inversion recovery (FLAIR), T2*-weighted imaging, and magnetic resonance angiography (MRA)^5^. Dynamic susceptibility contrast perfusion-weighted imaging (PWI) is often added to assess the ischemic penumbra, which is a key factor in determining eligibility for reperfusion therapies beyond standard time windows ^6-8^. PWI may also be instrumental in future efforts to identify candidates for recanalization in cases of distal occlusions or extensive infarcts beyond the traditional six-hour window^9^. However, PWI burdens the imaging workflow with increased costs and operational complexity by requiring contrast injection, longer scan time, and proprietary post-processing for parametric maps generation, such as time-to-maximum (T-max).

Recent advances in artificial intelligence (AI), in particular machine and deep learning, have opened up promising avenues for overcoming these limitations in emergency neuroradiology^10^. Over the past decade, research in medical image generation has made significant progress, with models ranging from convolutional U-Nets to more complex generative adversarial networks (GANs)^11^. Notably, GAN-based approaches have demonstrated the ability to synthesize FLAIR images from DWI^12^ and T2* from DWI^13^, suggesting the potential for abbreviated MRI protocols in which certain sequences are omitted during acquisition and reconstructed offline within seconds. Preliminary studies have also shown that perfusion maps may be inferred from MRA data alone, although the spatial congruence between synthetic and actual hypoperfused regions remains moderate^14^.

Building on these developments, our study aimed to provide a new perspective and go beyond the current state-of-the-art of image-to-image translation for stroke imaging. We hypothesized that together DWI and FLAIR might contain sufficient information to accurately estimate T-Max maps. For instance, a gradient of subtle diffusion restriction has been observed around the infarct core, with the possibility of defining penumbra based on mild apparent diffusion coefficient (ADC) decrease instead of PWI^15^. Additionally, FLAIR vascular hyperintensities extending beyond the DWI lesions have been associated with the presence of PWI-defined penumbra^16, 17^. Hence, generating T-max maps derived from DWI and FLAIR inputs appears feasible and may obviate the need for dynamic contrast-enhanced imaging in selected patients. Accordingly, newer and more robust architectures, such as diffusion models, may offer stability and generalizability for this purpose.

In this study, we anticipated that newer and more robust architectures, such as diffusion models, may offer stability and generalizability for this purpose. Therefore, we conducted a comprehensive evaluation of several diffusion-based models alongside the competitive GAN architecture^11^ to identify the optimal configuration for generating accurate T-max maps at the acute stage of stroke, using only non-contrast DWI and FLAIR as inputs, with the future perspective to enhance timely access to decision-critical data.

## Materials and Methods

This study is reported in accordance with the TRIPOD statement for prediction model development and validation studies.

### Patients

This study included imaging data from 355 patients with acute ischemic stroke sourced from two independent cohorts in Germany (Heidelberg, n=201) and France (Bordeaux, n=154). The Heidelberg data were used for training and validation, randomly split in an 80:20 ratio.

The Bordeaux data served for external validation, as these patients originated from a different country, healthcare system, and had been assembled independently of the Heidelberg cohort. The study was approved by the respective ethics committees. As it involved retrospective analysis of anonymized data collected as part of routine care, patients were informed of their participation in the study and given the option to withdraw.

Consecutive patients referred to the two centers for acute stroke were retrospectively included. In both centers, patients were identified from the hospital data warehouse using the following criteria: (i) confirmed ischemic stroke due to occlusion of the internal carotid artery terminus or of the M1/M2 segments of the middle cerebral artery, and (ii) MRI including PWI performed at the acute stage. We excluded the patients with ischemic stroke in another vascular territory, hemorrhagic stroke, or with MRI data of insufficient quality.

Demographic and clinical data were collected, including age, sex, vascular risk factors, acute NIHSS score, time from symptom onset to MRI, and acute treatments.

### MRI acquisitions and image pre-processing

MRI examinations from Heidelberg were performed on 3T scanners (Magnetom Verio, TIM Trio, and Magnetom Prisma; Siemens Healthcare, Erlangen, Germany) while those from Bordeaux were acquired using a 1.5T scanner (Magnetom Aera; Siemens Healthcare, Erlangen, Germany). Typical imaging parameters for DWI included a repetition time (TR) of 500 ms, echo time (TE) of 200 ms, a slice thickness of 5 mm, a field of view (FOV) of 500 × 500 mm, and a matrix size of 128 × 128, with b-values of 0 and 1000 s/mm^2^. FLAIR images were generally acquired with a TR of 9000 ms, TE of 120 ms, slice thickness of 5 mm, FOV of 230 × 230 mm, and matrix size of 256 × 256. PWI sequences used TRs of approximately 2200 ms, TEs around 30–36 ms, a 5 mm slice thickness, a 240 × 240 mm FOV, and a 128 × 128 matrix, with a gadolinium dose of 0.1 mmol/kg and an injection rate of 5 mL/s. These parameters were not strictly identical across cohorts, reflecting institutional protocol differences. To minimize the impact of inter-site variability, images were spatially co-registered, resampled to standardized dimensions, and intensity-normalized prior to model training (see below).

The dynamic contrast-enhanced images were converted to parametric time-to-maximum (T-max) maps using Olea Sphere® (Olea Medical, La Ciotat, France) based on deconvolution of the regional concentration-time curves with an automatically detected arterial input function. T-max masks were automatically generated by applying a threshold of >6 seconds, in accordance with established standards^6, 8^. Infarct core regions were delineated from DWI using an automatic lesion segmentation algorithm (ISLES 2022)^18^ for the Heidelberg cohort, and Olea Sphere for the Bordeaux cohort. This difference reflects site-specific clinical workflows. All DWI and PWI-derived masks were reviewed and, when necessary, manually refined by a junior radiologist (CM) and a senior radiologist (TT).

Raw DWI and FLAIR images, along with the corresponding T-max maps, underwent skull stripping and co-registration to native space using FSL, with FLAIR serving as the reference. Subjects with missing modality data (e.g., absent FLAIR, DWI, or T-max maps) were excluded from further analysis to ensure consistent preprocessing and spatial alignment. The resulting spatial transformations were applied to both DWI and PWI masks to ensure alignment.

For model training, each case was resampled to a slice thickness of 5.5 mm resulting in standardized and harmonized image dimensions (256 × 192 x 18 voxels) across samples to ensure consistency and compatibility with the deep learning pipelines.

### Methodological approaches for image synthesis

We employed a specialized type of deep learning models known as denoising diffusion probabilistic models (DDPM). DDPMs simulate a Markovian forward process by progressively adding Gaussian noise to input data, and then learn to reverse this process to generate high-quality images. Recently, diffusion models have gained significant attraction due to their superior performance in image synthesis compared to alternative approaches such as GANs.

To select the best model architecture, we experimented with different input channel configurations and input dimensionalities. We compared different models by varying the input dimensionality; a 2D model with one axial slice as input, 2.5 D with 7 consecutive slices as input (e.g., the maximum number of consecutive slices our computational resources allowed for), and a 3D model that accepted 3D volumetric inputs. Subsequently, to enhance prediction accuracy, we introduced a perfusion-weighted loss function. Specifically, the standard voxel-wise reconstruction loss was modified by applying increased weighting to voxels belonging to the ground truth T-max >6s mask. This weighting scheme penalizes errors more strongly within hypoperfused areas while preserving global image fidelity. The objective was to encourage the model to focus on spatially and clinically meaningful perfusion delays without compromising overall structural reconstruction. Both the 2.5D and 3D models were re-trained upon introducing this loss function modification. Lastly, we enhanced the 2.5D model, which yielded better results in the previous step by adding the binary infarct core mask derived from the DWI as a third input channel. These efforts resulted in the development of six distinct diffusion models in total for predicting T-max maps from the combination of FLAIR and DWI in 2D, 2.5D, and 3D, then 2.5D and 3D with a modified loss function, and for the latter with the addition of the infarct core mask.

Given that most cross-modality image synthesis studies rely on GAN-based architectures^11^, we also trained a model for a ‘state-of-the-art’ comparison with our diffusion models. Specifically, we implemented a 3D adaptation of the pix2pix GAN^19^, consisting of two networks: a generator with a U-Net-based architecture and a convolutional discriminator. The generator was tasked with synthesizing a 3D T-max map from concatenated DWI and FLAIR inputs, while the discriminator learned to distinguish between the generated data and the real T-max map. Implementation details of the GAN can be found in the referenced work in^14^. A graphical overview of the workflow with detailed steps is shown in Supplementary Figure 1 (S1).

### Training

Using 80% of the Heidelberg dataset, we trained a total of seven distinct models (6 DDPM + 1 GAN model), each designed to generate the perfusion parameter map T-max. To ensure a fair comparison across the different architectures and input combinations, all models were trained for 200 epochs. The default batch size was set to 16; however, for the more computationally demanding configurations, particularly the 3D models and the GAN, the batch size had to be reduced to 1 due to GPU memory constraints. Optimization was performed using the Adam optimizer with β_1_ = 0.9 and β_2_ = 0.95, and a constant learning rate of 1×10^-^□applied uniformly across all diffusion models. All training procedures were conducted on a NVIDIA Titan RTX GPU (NVIDIA Corporation, Santa Clara, CA, USA), and model development was implemented using the PyTorch framework.

### Performance Evaluation

Model performance was first evaluated on the internal validation set, comprising 20% of the Heidelberg dataset, followed by a comprehensive assessment on the external Bordeaux validation dataset. After visual inspection, the predicted T-max maps were refined by applying a patient-specific binary FLAIR mask to remove cerebrospinal fluid. Quantitative evaluation was based on four widely used image similarity metrics: the Structural Similarity Index Measure (SSIM), Peak Signal-to-Noise Ratio (PSNR), Pearson Correlation Coefficient (PCC), and Mean Absolute Error (MAE). SSIM captures differences in luminance, contrast, and structural information between predicted and ground truth images, while PSNR measures the ratio between the maximum possible signal and the noise. PCC assesses the linear correlation between voxel intensities in the predicted and true T-max maps. Higher SSIM, PSNR, and PCC indicate better similarity, whereas lower MAE values, calculated as the L1 norm, reflect higher predictive accuracy.

To assess the clinical relevance of the model outputs, we conducted an additional segmentation-based overlap analysis. Specifically, we assessed linear correlations between the predicted and ground truth binary PWI masks of T-max > 6 seconds and then computed the Dice coefficient to evaluate the spatial overlap, which ranges from 0 (no overlap) to 1 (perfect overlap). Binary masks from the synthetic T-max were generated by applying a threshold of T-max > 6 seconds. To eliminate non-relevant regions after the thresholding step and improve spatial continuity, we applied Gaussian smoothing with a sigma of 1.2 to the binarized predictions. The same processing was applied to the ground truth masks, which were generated using the OLEA software® and manually reviewed by two raters (TT and CM).

To further explore clinical applicability, we also examined the PWI-DWI mismatch, as a standard marker of penumbra. This analysis aimed to analyze the added value of synthetic perfusion images beyond diffusion imaging. Mismatch distributions were compared between synthetic and ground truth maps.

Finally, we investigated the contribution of each input modality to the overall model performance. Using the best-performing model as a baseline, we generated three additional sets of predictions, each omitting one of the input channels: FLAIR, DWI, or the infarct core mask. The outputs were evaluated using the same metrics, and the performance degradation was interpreted as an estimate of each modality’s relative importance in guiding accurate T-max map generation. This analysis was carried out using the Z-score normalization to quantify the deviation from the ground truth in terms of standard deviations.

## Results

### Patients

A total of 355 stroke patients who underwent acute MRI were included and divided into 160 from training, 41 for internal testing, and 154 for external validation. On average, patients showed large volumes of hypoperfusion based on T-max>6s as measured by PWI, with substantial variability reflecting differences in individual resistance to ischemia. Population characteristics can be found in Table 1.

**Table 1.** Patient Characteristics for the training, testing, and external validation datasets.

| Characteristics | training (Heidelberg)<br>(n=160) | testing (Heidelberg)<br>(n=41) | validation (Bordeaux)<br>(n=154) | p values† |
| --- | --- | --- | --- | --- |
| Female sex | 88 (55.0%) | 24 (58.54%) | 92 (60.0%) | ns |
| Age (years) | 67.81 ± 14.89 | 68.34 ± 12.92 | 75.5 ± 13.7 | <0.0001 |
| **Vascular risk factors and medical history** |  |  |  |  |
| Hypertension | 100 (62.5%) | 24 (58.54%) | 95 (61.5%) | ns |
| Hyperlipidemia | 44 (27.5%) | 12 (29.3%) | 54 (35.1%) | ns |
| Diabetes | 29 (18.12%) | 7 (17.07%) | 23 (15.0%) | ns |
| Active smoking | NA | NA | 26 (17.0%) | NA |
| Previous Stroke | NA | NA | 16 (10.5%) | NA |
| Myocardial infarction | 28 (17.5%) | 7 (17.07%) | 17 (11.0%) | ns |
| Atrial fibrillation | 55 (34.38%) | 18 (43.90%) | 30 (19.5%) | 0.0002 |
| **Stroke characteristics and acute treatment** |  |  |  |  |
| Baseline NIHSS score | 15.57 (0–33) | 16.32 (2–25) | 14 (7–19) | 0.0015 |
| Time from symptom onset to MRI (minutes) | 419.52 (29– 5123) | 370 (14–1225) | 239.5 (143–455) | 0.033 |
| Infarct core volume on DWI (mL) | 21.27 ± 18.58 | 22.41 ± 23.25 | 22.17 ± 26.93 | ns |
| **Volume of T-max>6s (mL)** | 93.48 ± 65.65<br>(0.79 mL– 395.96 mL) | 108.56 ± 77.50<br>(0.67 mL – 312.71 mL) | 103.76 ± 90.98<br>(0.00 mL – 605.01 mL) | ns |
| **T carotid occlusion** | 25 (15.62%) | 5 (12.20%) | 15 (9.5%) | ns |
| **M1 occlusion** | 100 (62.50%) | 28 (68.29%) | 96 (62.5%) | ns |
| **M2 occlusion** | 27 (16.88%) | 7 (17.07%) | 58 (37.5%) | 0.004 |
| **IV thrombolysis** | 97 (60.62%) | 27 (65.85%) | 91 (59.5%) | ns |
| **Mechanical thrombectomy** | 160 (100%) | 41 (100%) | 93 (60.5%) | <0.0001 |

Data are numbers (%), mean ± SD or median (Q1-Q3)
† Comparisons are between Heidelberg (full cohort) and Bordeaux using chi<sup>2</sup>, t-test or Mann-Whitney as appropriate.
\* Due to missing data, the stroke characteristics (occlusion type) columns for the Heidelberg dataset do not add up to 100%, whereas in the Bordeaux dataset, multiple occlusions were reported for some patients, making the sum exceed 100%.

### Toward the best-performing model

Through a stepwise increase in model dimensionality and complexity, we trained, evaluated, and compared seven distinct models (6 diffusion models + 1 GAN model, Figure S1). We began with a 2D diffusion model, where both inputs and outputs consisted of single 2D image slices. Although individual slice predictions were promising, reconstructing them into a full 3D volume revealed limitations in capturing inter-slice dependencies. The resulting volumes appeared fragmented and lacked smooth anatomical continuity (Figure 1).

To overcome this, we developed a 2.5D model that processed input data in overlapping chunks of seven slices. This approach better retained inter-slice contextual information and produced more coherent 3D reconstructed outputs.

**Figure 1.**
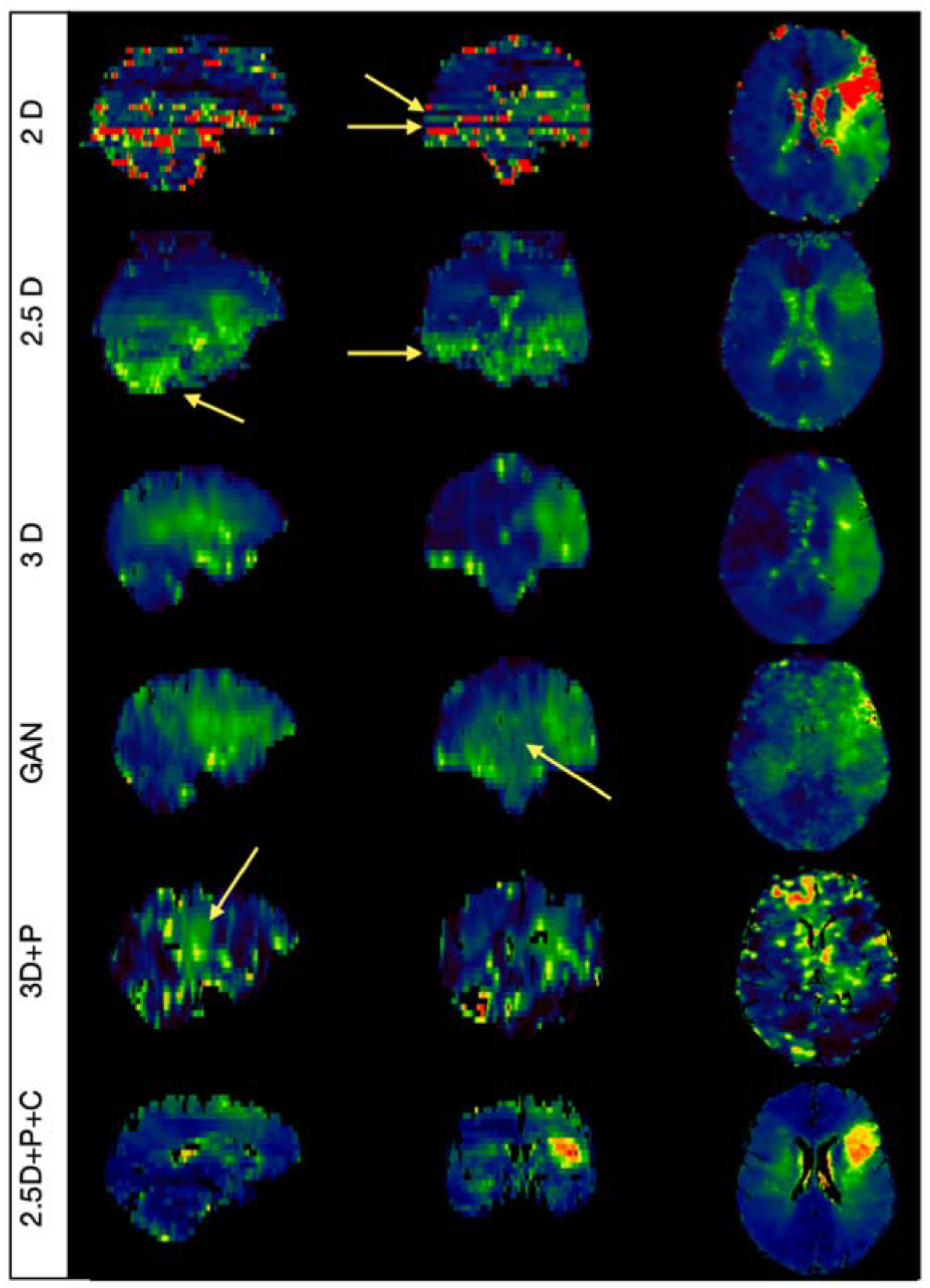
Overview of the approaches taken toward the best-performing model. Illustrative synthetic T-max maps are shown in sagittal, coronal and axial planes. Discussed details of the occurring problems are indicated by the yellow arrows, which point towards e.g., scattered reconstruction in the 2D model; overestimation in the cerebellum for the 2.5D model, and instabilities in the predictions for the 3D models and the GAN.

Despite this improvement, the model tended to replicate and propagate artifacts present in the ground truth data, particularly in regions near the cerebellum and temporal lobes (Figure 1). To address this, we introduced a modified loss function incorporating binary perfusion masks, referred to as the ‘perfusion loss’ (see Methods). This adjustment served two key purposes: it strengthened the model’s capacity to learn spatial dependencies across slices, and it emphasized accurate intensity predictions in hypoperfused areas by penalizing errors more heavily in those areas.

We also trained a full 3D model, both with and without the perfusion-weighted loss function. As expected, the limited training dataset (comprising 480 images from 160 patients, including the DWI, FLAIR, and Infarct Core mask for each patient) proved inadequate for robust 3D learning, resulting in poor generalization.

In parallel, we implemented a 3D pix2pix GAN as a state-of-the-art comparator^11^. While this model demonstrated intermittent success by unreliably yielding accurate results, its training was unstable, likely reflecting the same data limitations. Given these outcomes and in light of the strong and consistent performance of the 2.5D architecture, we selected this intermediate-dimensional model as the most promising architecture.

As a final enhancement, we incorporated as an additional input channel, the automatically derived binary infarct core mask from DWI. This mask offered valuable additional spatial context, helping the model differentiate infarcted from at-risk tissue. This modification further improved visual predictive performance, establishing the 2.5D model with both perfusion loss and infarct core mask input (2.5D+P+C) as the final and most promising model for application on both internal and external validation. An overview of the seven models tested, alongside their performance is visualized in Figure S2.

For the selected 2.5D+P+C model, the generation of the predicted chunks and the reconstruction of one complete 3D T-max map (sampling) required on average 1,78 minutes (107,37 seconds) using a single NVIDIA TITAN RTX GPU.

### Validation testing set

Model performance was validated through quantitative analysis on an independent test subset, comprising 20% of the Heidelberg dataset, held out from the start of the study.

Among all models, the 2.5D+P+C configuration consistently achieved the highest performance, with a mean structural similarity index (SSIM) exceeding 0.82, a peak signal-to-noise ratio (PSNR) above 20.96, and a mean absolute error (MAE) below 0.032. Voxel-wise intensity correlations with the ground truth yielded a Pearson correlation coefficient of r = 0.65 (Table 2).

**Table 2.** Overview of the quantitative analyses for the models’ internal validation (n=41). In the model descriptions, P stands for perfusion loss and C for the addition of the infarct core mask. SSMI: mean structural similarity index; PSNR: peak signal-to-noise ratio; MAE: mean absolute error; PCC: Pearson Correlation Coefficient. MVD: Mean Volumetric Difference. Data are mean (standard deviation). For each metric, the bold values indicate the best result. The quantitative analyses for the 2.5D+P+C external validation (n=154) are also indicated.

|  | SSMI | PSNR | MAE | PCC | DICE | MVD (mL) |
| --- | --- | --- | --- | --- | --- | --- |
| **Out-of-sample INTERNAL validation (N=41)** |  |  |  |  |  |  |
| **2D** | 0.786 (0.18) | 17.83 (6.69) | 0.040 (0.02) | 0.386 (0.06) | 0.503 (0.15) | 68.73 (54.1) |
| **2.5D** | 0.796 (0.18) | 18.10 (7.52) | 0.038 (0.01) | 0.590 (0.08) | 0.352 (0.11) | 123.49 (88.2) |
| **2.5D+P** | 0.782 (0.18) | 17.55 (7.67) | 0.043 (0.01) | 0.569 (0.08) | 0.241 (0.10) | 336.11 (128.5) |
| **2.5D+P+C** | **0.820 (0.03)** | **20.96 (4.29)** | **0.032 (0.01)** | **0.650 (0.09)** | **0.802 (0.08)** | **21.44 (18.9)** |
| **3D** | 0.788 (0.16) | 18.59 (6.64) | 0.038 (0.01) | 0.407 (0.07) | 0.412 (0.16) | 72.71 (55.9) |
| **3D+P** | 0.793 (0.17) | 18.27 (6.98) | 0.039 (0.01) | 0.466 (0.09) | 0.338 (0.13) | 79.57 (77.1) |
| **GAN** | 0.802 (0.18) | 19.27 (7.12) | 0.035 (0.03) | 0.603 (0.10) | 0.411 (0.13) | 96.99 (75.4) |
| **Out-of-sample EXTERNAL validation (N=154)** |  |  |  |  |  |  |
| **2.5D+P+C** | 0.863 (0.02) | 25.29 (12.56) | 0.022 (0.01) | 0.641 (0.08) | 0.597 (0.13) | 55.88 (54.3) |

To assess clinical relevance, we further analyzed volumes of significant hypoperfusion, defined by T-max > 6 seconds. Volumetric comparisons between binary masks derived from synthetic and ground truth T-max maps showed particularly high agreement for the final 2.5D+P+C model (r = 0.935), as visualized in Figure 2A. This strong correlation was accompanied by the highest mean Dice coefficient (0.802), indicating robust spatial overlap (Table 2). Notably, correlation remained stable across small lesion volumes for the 2.5D+P+C model, in contrast to other configurations. This likely reflects the combined effect of perfusion-weighted loss and infarct core guidance, which might improve spatial consistency and reduce false-positive predictions in low-volume cases.

**Figure 2.**
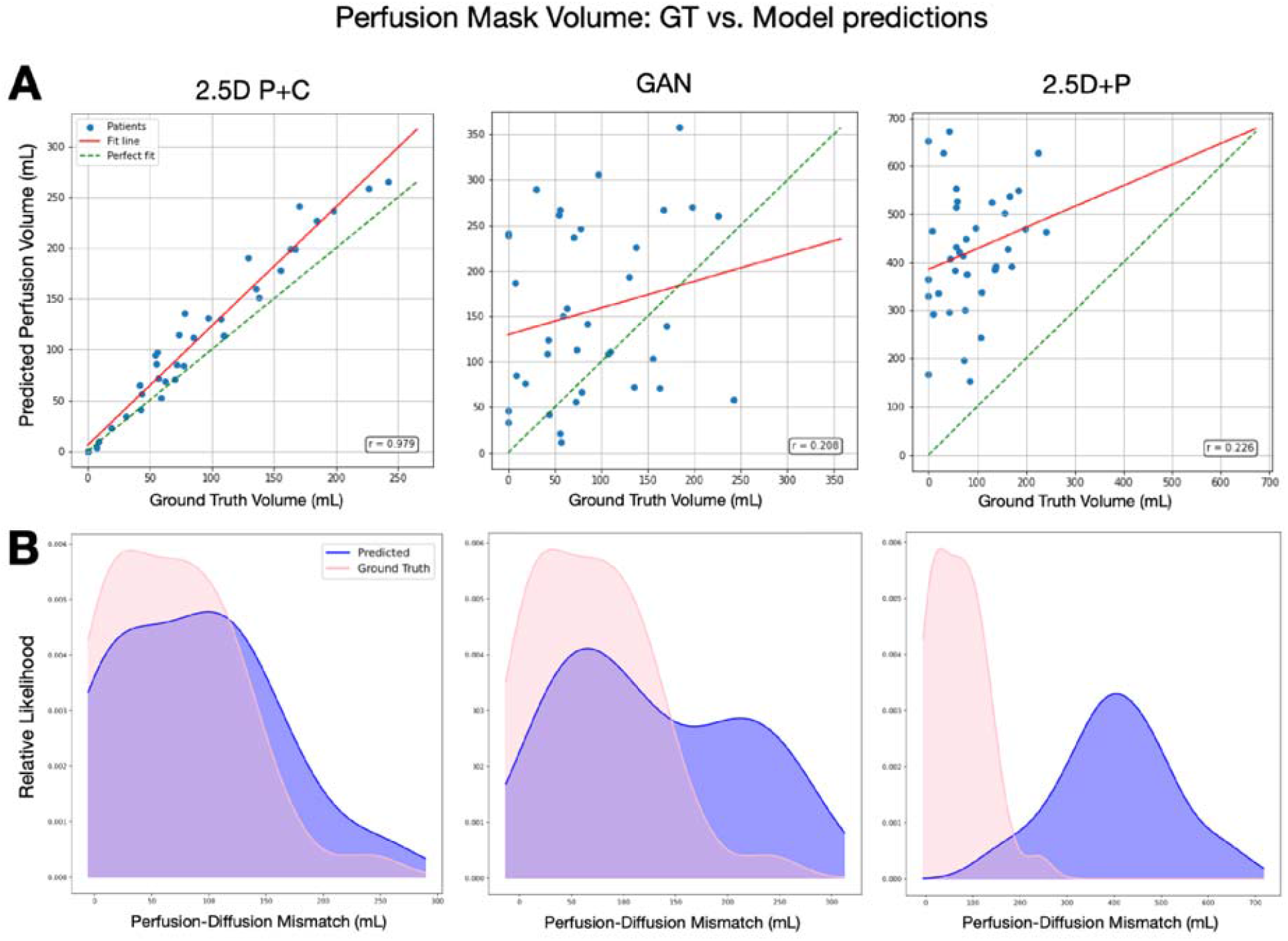
**A** Scatter plot of perfusion mask volumes, ground truth volumes plotted against the perfusion mask derived from the predicted models (the three best performers are depicted). The red line represents the fitted line for the model, whereas the green line represents a perfect fit. **B** Distributions of the relative likelihood of the perfusion-diffusion mismatches for the same modes. The pink curves represent the mismatch distribution for the ground truth data, while the purple curves represent the mismatch distributions for the predicted models.

While the model effectively localized hypoperfused regions, intensity values tended to be mildly underestimated. After applying a correction based on the mean predicted intensity, we recalculated perfusion–diffusion mismatch volumes. The adjusted outputs again showed the closest alignment with the ground truth for the 2.5D+P+C model, confirming its superior fidelity in capturing clinically meaningful perfusion patterns (see Figure 2B). Figure 3 shows one illustrative example of the synthetic perfusion from the 2.5D+P+C for a typical patient with a large volume of mismatch.

**Figure 3.**
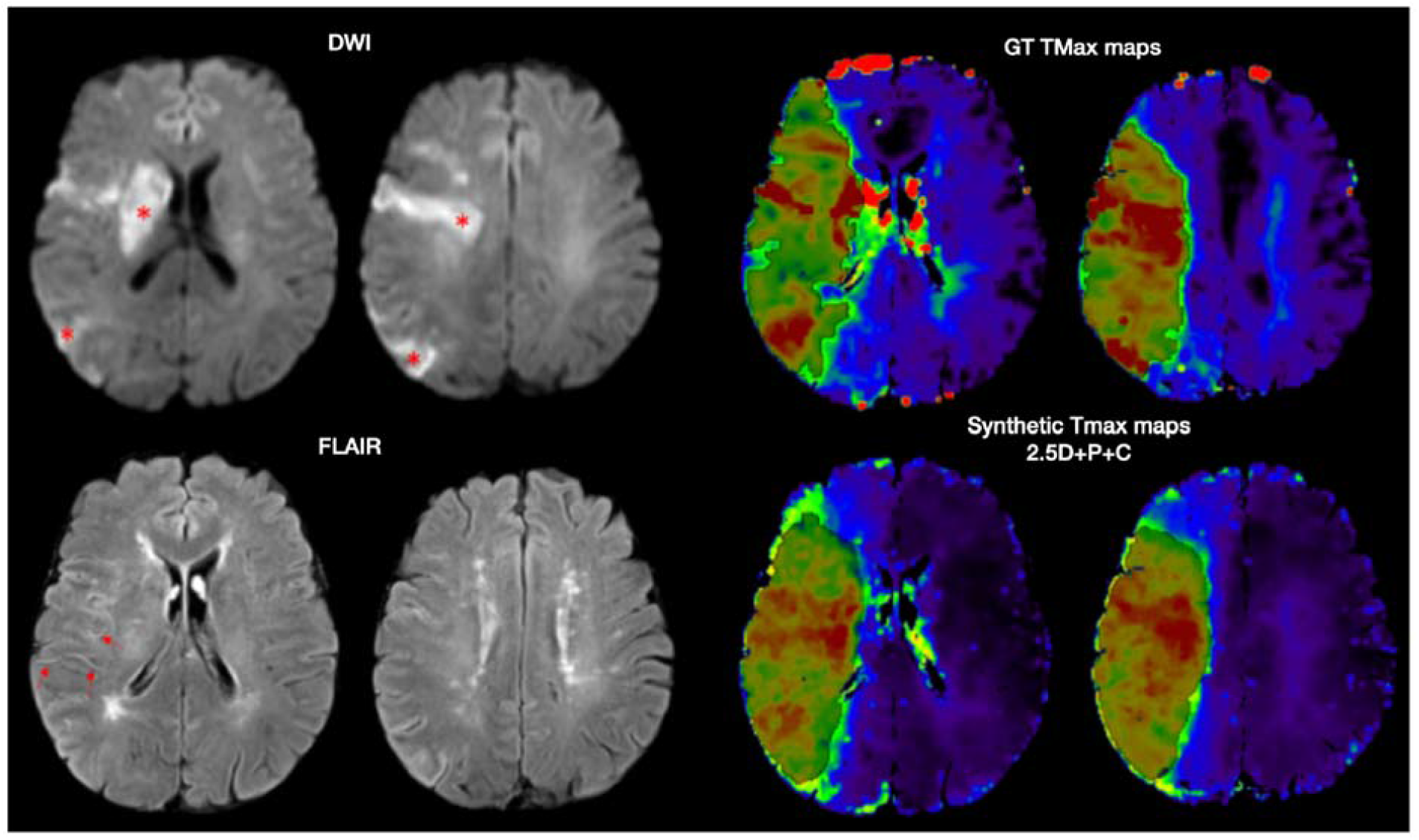
Illustrative example of a 69-year-old patient with a right acute stroke explored 216 minutes from the symptom onset. The DWI shows the infarct core (*), possibly associated with more subtle surrounded diffusion restriction and with FLAIR vascular hyperintensities (arrows) aligning with a large amount of penumbra (T-max>6s was outlined in black) that has been closely captured with the 2.5D+P+C synthetic map (Dice=0.805).

### Relative weight of each modality

To better understand the contribution of each input modality, we conducted an ablation analysis using the final 2.5D+P+C model. We generated three alternative sets of predictions, each omitting one input: FLAIR, DWI, or the Infarct core mask, and evaluated them using the same metrics (Figure 4A). Performance dropped most significantly when the infarct core mask was excluded, with the average Dice coefficient falling from 0.802 (full model) to 0.476. Exclusion of DWI and FLAIR resulted in smaller, though still meaningful declines (average Dice scores of 0.609 and 0.676, respectively). This pattern was further corroborated by mean Z-score analyses (Figure 4B), which quantify the deviation from the ground truth: FLAIR yielded a Z-score of 0.115 (SD = 0.74), DWI a Z-score of 0.137 (SD = 0.79), and the infarct core mask a notably higher Z-score of 0.241 (SD = 0.998). These findings indicate a moderate and relatively stable effect of excluding DWI or FLAIR, but a larger and more variable impact when the infarct core mask was excluded. Further analysis revealed a positive correlation between core volume and Z-score, indicating that the predictive importance of the infarct core mask increases with infarct core size. This relationship, illustrated in Figure 4C, highlights the critical role of the infarct core in guiding accurate perfusion map synthesis. The model was, however, able to predict perfusion modifications outside the visible DWI core as shown above with the mismatch analysis.

**Figure 4.**
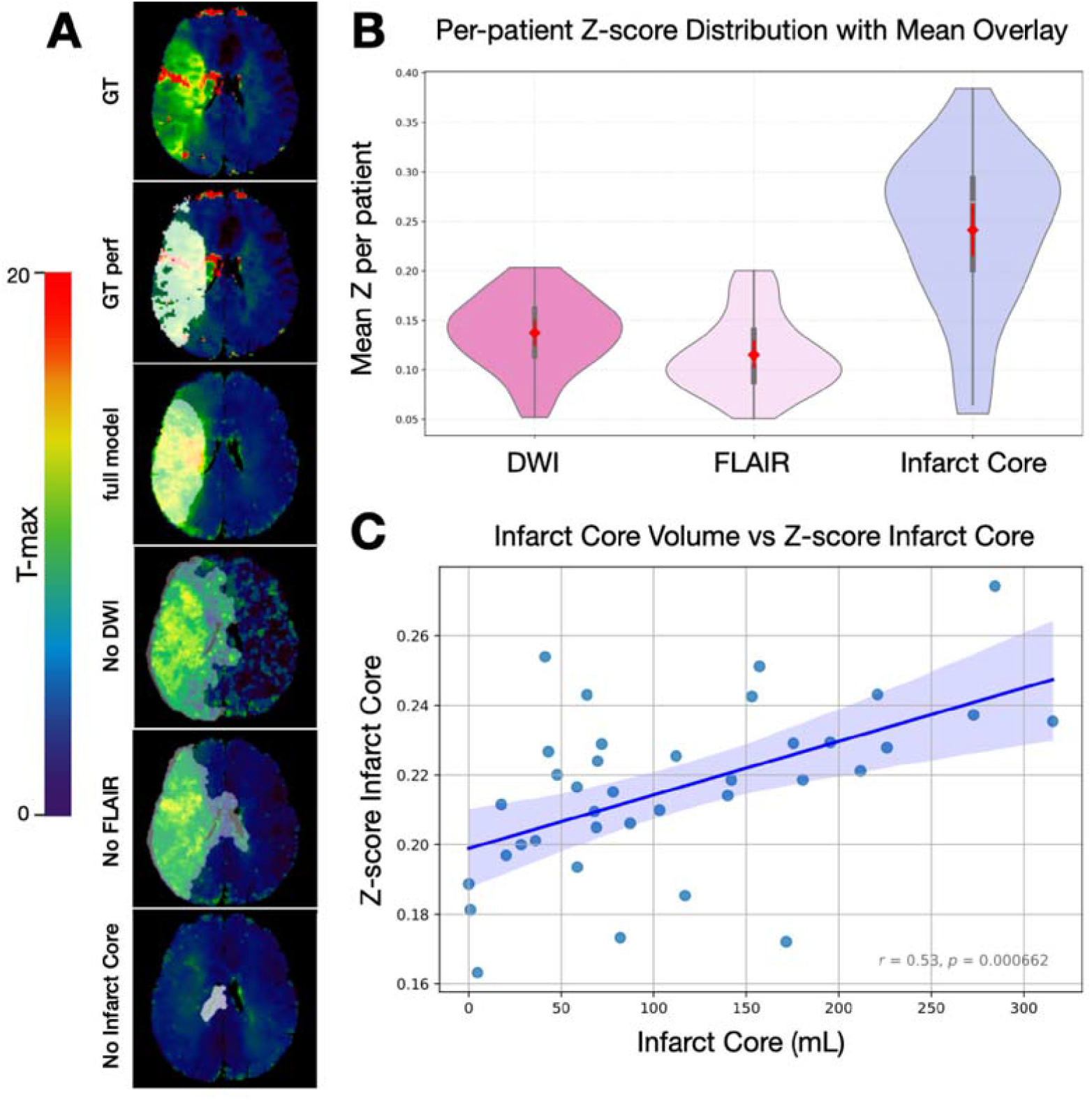
Assessment of input modality contributions to synthetic perfusion map accuracy. **A** Visual demonstration of predictions with the final 2.5D+P+C model upon omitting each of the inputs (DWI, FLAIR, Infarct Core). The first two rows show the ground truth (GT) T-max maps and the highlighted GT perfusion mask, respectively. The third image from the top represents the output of the full model, which contains all inputs to generate a synthetic T-max map. The semi-transparent (white) highlighted areas indicate the resulting perfusion mask based on the model outputs (T-max>6). **B** Violine plots of the Z-score analysis showing the effect of omitting DWI, FLAIR, and infarct core, as model input, respectively. Each violin represents the distribution across patients, with the red diamond indicating the group mean and the vertical red line denoting the standard error of the mean. The plot demonstrates higher mean Z-scores and variability for the core data compared to DWI and FLAIR modalities. **C** Scatter plot showcasing a positive correlation between the core volume and the Z-score.

### External validation

External validation was conducted on a fully independent cohort comprising 154 patients from the Bordeaux dataset, none of whom were used during model training or hyperparameter tuning. The final 2.5D+P+C model was assessed using the same evaluation protocol as for internal validation.

As expected, performance metrics were lower compared to the internal validation set, reflecting inherent differences in patient demographics and MRI acquisition parameters across sites. Nevertheless, the model maintained good generalizability, achieving a mean Dice coefficient of ≈ 0.6, which still indicates clinically meaningful spatial overlap.

Interestingly, average image similarity metrics (e.g., SSIM and PSNR) slightly surpassed those observed in internal validation (Table 2). This trend may be attributed to the larger sample size and potential smoothing effects related to inter-individual variability. Two representative examples from this cohort (for a successful and an unsuccessful output) are shown in Figure S3.

## Data availability

The datasets used in this study are not publicly available. They are, however, available from the corresponding author upon reasonable request and subject to institutional and regulatory approval where required.

## Discussion

Our study demonstrates that a deep learning framework can reliably generate synthetic T-max perfusion maps from non-contrast DWI and FLAIR clinical sequences. This approach may offer a compelling alternative to dynamic susceptibility contrast PWI, with the potential to streamline acute stroke imaging, reduce delays in decision-making, and expand access to perfusion data without the need for contrast administration or specialized post-processing.

In acute stroke care, where “time is brain,” efficiency is crucial. While MRI-based protocols are used in some comprehensive stroke centers, the typical median scan time is 13 minutes^4^. Although acceleration techniques have improved acquisition efficiency^5, 20^, dynamic susceptibility contrast (DSC) perfusion remains limited by the temporal requirements of capturing the complete first-pass of contrast. This is typically locked into a 1.5 to 2-minute window and cannot be shortened, as it depends on physiological variables such as cardiac output, injection timing, and vascular status. In practice, total time consumption is even greater due to contrast preparation, inter-sequence delays, and the post-processing required to generate parametric maps from raw data. Real-world evidence shows that omitting DSC-PWI can reduce MRI duration from a median of 16 minutes (IQR, 15–19) to 12 minutes (IQR, 10– 15)^21^. However, this reduction comes at the cost of forfeiting valuable perfusion information. Arterial spin labeling (ASL) has also been proposed as a non-contrast alternative for assessing cerebral blood flow and tissue at risk^22^ but is limited by lower signal-to-noise ratio, longer acquisition times, and sensitivity to motion and delayed transit effects^22^. Our approach addresses these limitations by enabling the inference of perfusion maps directly from standard, non-contrast sequences (DWI and FLAIR), and thereby eliminating the need for direct DSC-PWI acquisition. Since these sequences are already part of routine stroke protocols, their use for generating synthetic perfusion maps incurs no additional scan time. Moreover, the reconstruction process can be initiated concurrently with the acquisition of other sequences such as T2* or MRA, potentially making perfusion data available in real time and “for free” in all patients.

Currently, PWI is primarily used to identify mismatch in patients considered for extended-window intravenous thrombolysis ^6, 7^ or late-window thrombectomy ^8, 23^. Although recent randomized trials have demonstrated benefit of thrombectomy in patients with large infarct cores without strict mismatch-based selection criteria ^24-28^, perfusion imaging remains incorporated in current international guidelines for extended-window thrombolysis and thrombectomy ^29, 30^. Moreover, perfusion imaging may continue to assist in treatment decisions for large-core populations and extended time window where the benefit is still under investigation ^9^, distal vessel occlusions ^31^, mild stroke severity ^32^, very late presenters ^33^, and in the stratification of patients for future neuroprotective trials ^34, 35^. In this evolving therapeutic landscape, synthetic perfusion mapping may provide a pragmatic, non-contrast surrogate of perfusion delay, potentially expanding access to penumbral information while maintaining workflow efficiency.

Previous research has explored the feasibility of image synthesis in stroke, particularly the generation of FLAIR and T2* from DWI^12, 13^. These studies leveraged the embedded information in b0 images to reconstruct clinically useful sequences, establishing a precedent for image-to-image translation in deep learning in acute stroke imaging. More recently, Lohrke et al.^14^ proposed generating perfusion maps from time-of-flight MRA using a pix2pix GAN architecture, reporting a mean Dice score of 0.48 for regions with T-max > 6 seconds in a small internal dataset. Our findings demonstrate a substantial improvement (Dice coefficients reaching 0.80 on internal validation and 0.60 on external validation). These gains likely stem from both architectural enhancements and the use of more informative input channels. Specifically, the adoption of denoising diffusion probabilistic models, known for their stability and generalizability, represents an advantage over traditional GANs in this present case^11, 36^. Among the input features, the inclusion of infarct core masks provided critical spatial and pathological context, allowing the model to better differentiate infarct core from surrounding penumbral tissue. This is consistent with prior work describing a gradient of ADC decline around the infarct core, which may serve as a surrogate marker for tissue at risk^21, 37, 38^. While earlier attempts to define ADC-based penumbra^15, 39^ were limited by overlapping values between normal and threatened tissue, deep learning offers a more data-driven, nuanced interpretation of these subtle variations. Importantly, our approach does not rely solely on vascular imaging. Although MRA is effective in identifying proximal occlusions and modeling downstream perfusion deficits, it often fails to account for collateral flow, arguably the most relevant determinant of penumbral viability. Even within identical occlusion types, T-max volumes demonstrated marked heterogeneity, indicating that occlusion site alone does not determine downstream hypoperfusion. Synthetic perfusion mapping from parenchymal sequences such as DWI and FLAIR, therefore, offers a complementary perspective, potentially capturing information that conventional angiographic may overlook.

Generalizability remains a key challenge in medical AI, as models trained on data from a single institution often exhibit performance degradation when applied to external datasets^40^. To rigorously assess our model’s robustness and real-world applicability, we withheld the Bordeaux dataset from training and used it exclusively for external validation. This approach enabled us to evaluate generalizability across naturally occurring variations in clinical imaging and stroke populations. As expected, performance metrics declined on external data compared to internal validation, but retained strong volumetric correlations and a Dice coefficient close to 0.6. The mean volumetric difference (approximately 56 mL externally versus 21 mL internally) likely reflects the combined effects of acquisition variability, differences in infarct core segmentation tools, broader lesion heterogeneity, and magnetic field strength variation. The moderate reduction in spatial overlap observed in the external cohort likely reflects, in part, the acquisition differences. Nevertheless, the preservation of overall image similarity metrics and volumetric correlations suggests that the model retained sensitivity to physiologically meaningful patterns. These findings indicate preserved performance across institutions, while also highlighting the importance of further validation and potential adaptation strategies prior to clinical deployment. In future work, generalizability could be further enhanced by training on larger, multi-center datasets and incorporating further additional harmonization techniques such as z-score normalization, histogram matching^41^, or domain adaptation strategies^42^, including GAN-based frameworks that translate image domains to mitigate inter-institutional variability^43^. However, in the present study, we opted to test whether the diffusion model could inherently learn domain-invariant representations in an end-to-end pipeline without explicit normalization. Furthermore, we prioritized a low-latency inference workflow to ensure real-time clinical feasibility. This minimally processed setup is thought to serve as a baseline for future studies aiming further improve cross-site performance and generalizability.

Diffusion models are gaining attention in medical imaging due to their stability, sample diversity, and probabilistic nature^44^. Unlike GANs, which rely on adversarial training and often suffer from mode collapse, diffusion models learn to iteratively reverse a noising process and produce more reliable and consistent outputs^44^. These properties make diffusion models particularly well-suited for tasks involving uncertainty modeling, such as the modality translation explored in this study. Here, they proved especially effective in synthesizing perfusion maps, showing greater robustness compared to a 3D pix2pix GAN baseline. However, these benefits come with significant computational demands^45^. Training requires large datasets, substantial memory resources, and high computational overhead, particularly for full 3D implementations. We addressed this by adopting a 2.5D training strategy, feeding overlapping slice stacks into the network. This approach preserved spatial context while increasing training samples, striking a balance between model expressiveness and feasibility. While this added computational load during training and inference, the model still generated synthetic maps in under 110 seconds on average, compatible with real-time workflows in emergency settings.

Nonetheless, limitations remain. Our model was trained exclusively on patients with confirmed ischemic stroke who underwent acute MRI with DSC-PWI, and the study population therefore represents a selected subgroup of patients undergoing MRI-based stroke workup rather than the full unselected acute stroke population at each center. Consequently, the model’s performance in stroke mimics remains unknown, although PWI can be useful for identifying mimics^46^, and synthetic perfusion maps trained without such examples may not reproduce this diagnostic value. In addition, patients unable to undergo MRI or not selected for perfusion imaging may differ systematically in clinical severity, contraindications, workflow characteristics, or treatment pathways, which may limit the generalizability of the model. Additionally, DSC-PWI provides broader hemodynamic information beyond delay-based parameters alone, including cerebral blood flow and cerebral blood volume metrics. Our framework is designed to approximate clinically actionable T-max delay information in confirmed ischemic stroke, and should not be interpreted as a full replacement for comprehensive perfusion imaging in diagnostically uncertain cases. Furthermore, while the model generates continuous T-max maps, clinical interpretation often relies on dichotomized thresholds (e.g., T-max > 6s) to guide treatment decisions^47^. In some cases, predicted values fell just below the threshold, reducing binary overlap metrics despite accurate spatial localization. Future models could be trained to predict both continuous and binary maps or optimized specifically for threshold-based classification. Finally, while infarct core masks were provided as model inputs, their automatic generation is already supported in many stroke workflows and continues to improve^18^. Thus, this requirement should not pose a barrier to real-world implementation.

## Conclusion

We developed a deep learning framework capable of synthesizing T-max perfusion maps from routinely acquired, non-contrast MRI DWI and FLAIR sequences in the context of acute ischemic stroke. Our approach addresses a critical clinical need by offering a non-invasive, time-efficient alternative to traditional perfusion imaging, thereby contributing to faster, more accessible decision-making in the acute phase of stroke. While opportunities remain to further improve cross-institutional generalizability, this work lays the foundation for integrating synthetic perfusion mapping into real-world clinical workflows.

## Non-standard abbreviations and acronyms

ADC: Apparent diffusion coefficient
AI: Artificial intelligence
ASL: Arterial spin labeling
DDPM: Denoising diffusion probabilistic model
DSC: Dynamic susceptibility contrast
DWI: Diffusion-weighted imaging
FLAIR: Fluid-attenuated inversion recovery
FOV: Field of view
GAN: Generative adversarial network
MAE: Mean absolute error
MRA: Magnetic resonance angiography
NIHSS: National Institutes of Health Stroke Scale
PCC: Pearson correlation coefficient
PSNR: Peak signal-to-noise ratio
PWI: Perfusion-weighted imaging
SSIM: Structural Similarity Index Measure
T-max: Time-to-maximum
TE: Echo time
TR: Repetition time

