## Supplementary Information for "Generating Synthetic MR Perfusion Maps from DWI and FLAIR in Acute Ischemic Stroke: Development and External Validation of a Deep Learning Model"

### SUPPLEMENTAL MATERIAL

#### Diffusion model setup

For the diffusion models, we adopted a U-Net architecture with Group Normalization (GroupNorm) configured to eight groups and a base channel size of 128. Temporal information was incorporated via sinusoidal time embeddings. The models were trained using V-space prediction, with a MinSNR loss weighting ($\gamma$ = 5) to enhance training stability. An $\alpha$-cosine noise schedule was employed to guide the diffusion process. To further stabilize sampling and prevent divergence, value clipping was applied after each denoising step.

#### Supplementary Figures:

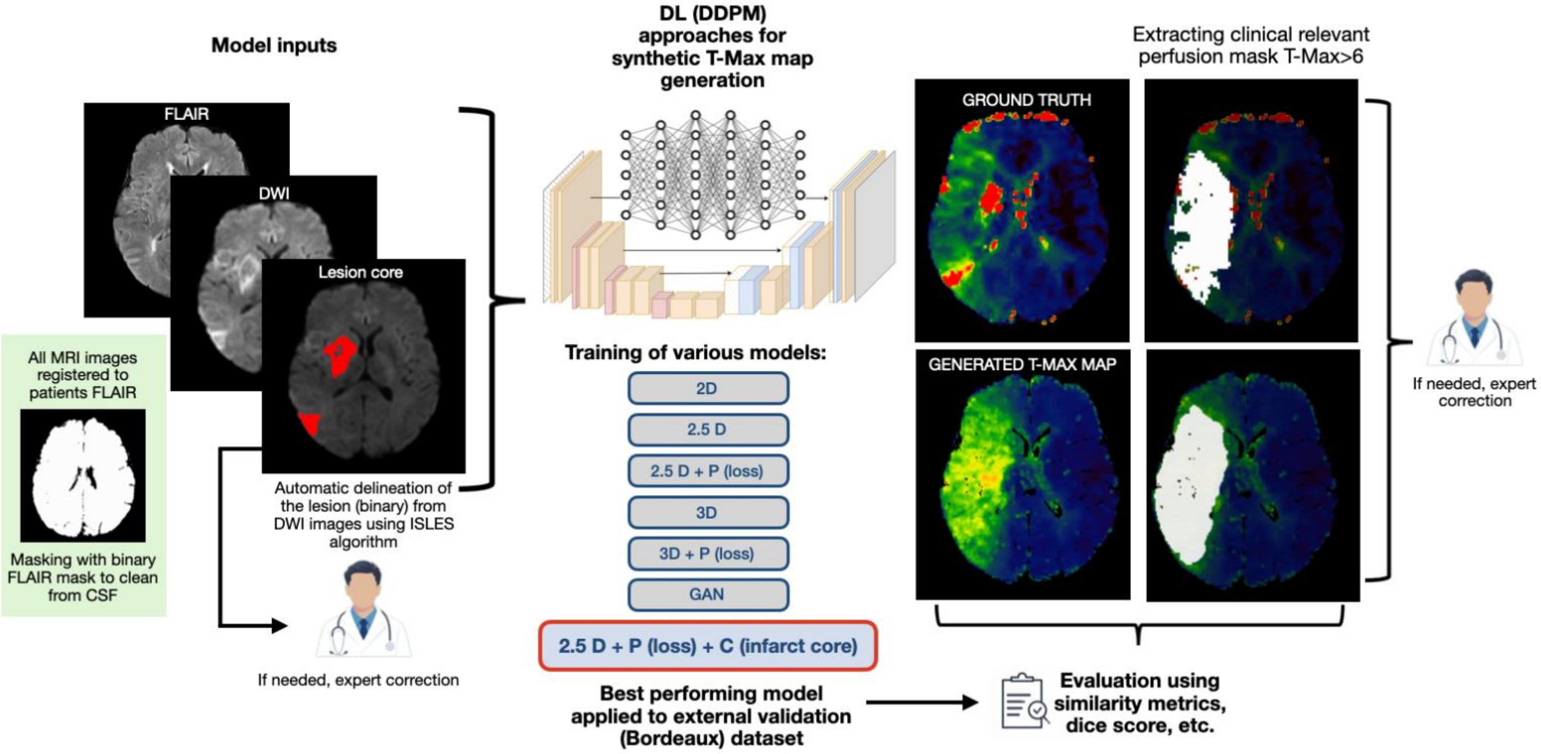


**Figure S1.** Workflow of the study visualized. Abbreviations: CSF+Cerebro Spinal Fluid; DL=Deep Learning; D=Dimension; P=modified perfusion loss function; C= infarct core; GAN=Generative Adversarial Network.

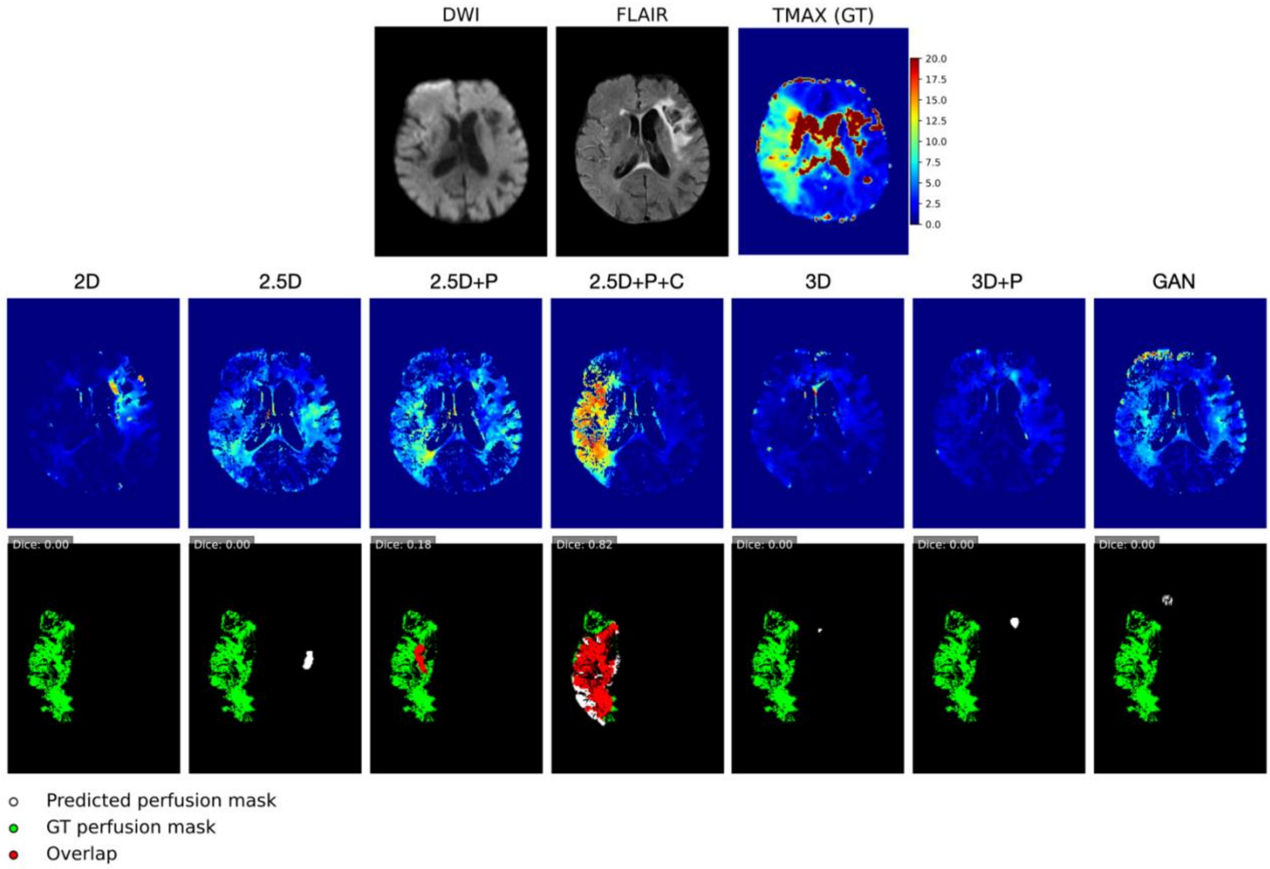


**Figure S2.** Overview of the seven different models that were tested, visualized on an exemplary patient's axial slice. The upper row shows the core inputs (DWI and FLAIR), as well as the ground truth (GT) T-Max map. The middle row shows the output T-max maps of each model for this particular slice. The bottom row represents the perfusion areas derived from the T-Max maps (>6 seconds), with the green mask marking the GT, the white area marking the predicted perfusion mask, and the red area marking the overlap between the ground truth and the predicted perfusion mask. The corresponding dice coefficient can be found in the upper left corner. (TMAX= time-to-maximum; GT=ground truth; P=perfusion loss; C=ischemic core, GAN=Generative Adversarial Network). Note: In this example, the patient has an old stroke (visible on FLAIR in the right hemisphere), which is correctly detected by the selected model (2.5D+P+C) as irrelevant, while less well-performing models wrongly include this information into their synthetic perfusion images.

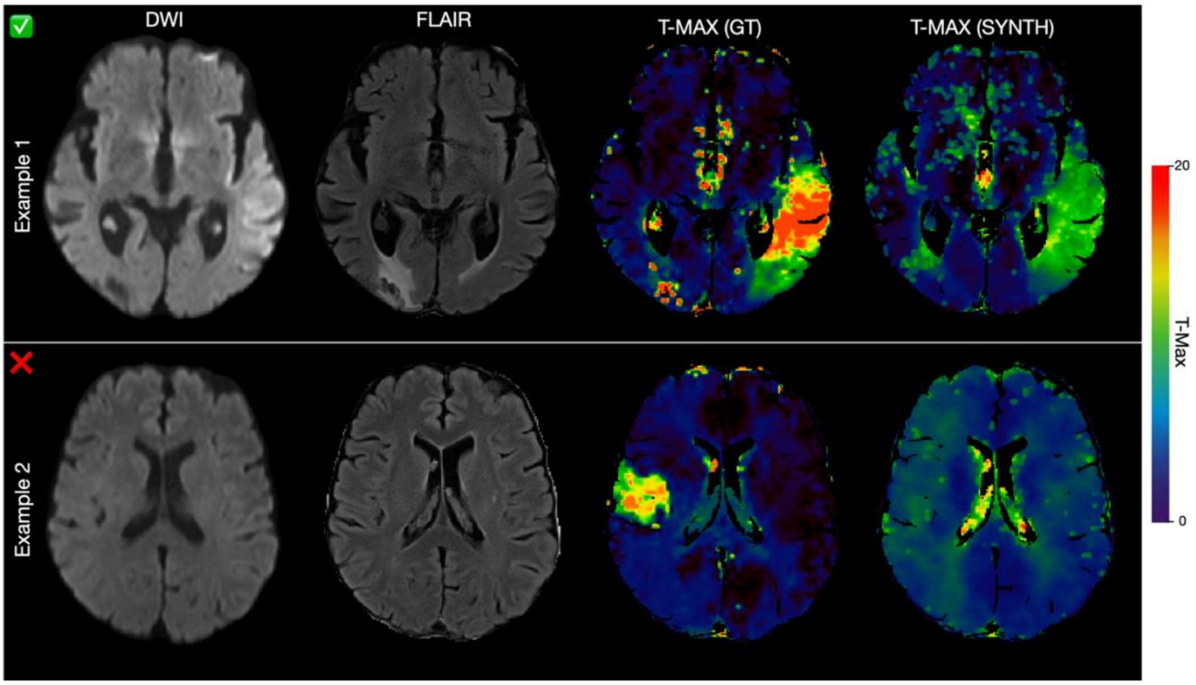


**Figure S3.** Two examples of outputs for the external validation using the final model (2.5D+P+C). Example 1 shows a successful generation of a synthetic T-Max map, whereas Example 2 demonstrates that the model can fail if the input images do not carry a sufficient amount of information to synthetically generate an accurate perfusion (T-Max) map.
